# Macrophages drive the inflammatory phase in experimental osteoarthritis

**DOI:** 10.1101/2020.05.28.122408

**Authors:** Anna Montgomery, Niamh Fahy, Samuel Hamilton, Blair Eckman, Lucia De Almeida, Shingo Ishihara, Maximilian G Mayr, Shang Yang Chen, Gaurav Gadhvi, Carla Cuda, Anne-Marie Malfait, Yvonne Bastiaansen-Jenniskens, Deborah R. Winter

## Abstract

Macrophages fulfill critical functions in maintaining tissue homeostasis in steady-state, as well as in inflammation and immune response. Inflammation is not considered a major driver of osteoarthritis (OA), but macrophages have been implicated in its pathogenesis. Here, we use two mouse models of experiment OA – collagenase-induced osteoarthritis (CIOA) and destabilization of the medial meniscus (DMM) – to quantify the immune cell infiltration into the knee joint during the early stages of disease. We find that the peak of inflammation occurs at day 3 in CIOA and is characterized by a transitory increase in neutrophils and monocytes and a longer-lived expansion of synovial macrophages. Macrophage sub-populations are disproportionally expanded with CX3CR1+ cells accounting for a larger proportion of the macrophage compartment. Transcriptional profiling demonstrates that synovial macrophages up-regulate inflammatory genes coinciding with peak inflammation and down-regulate genes associated with homeostasis and tissue-residence. Female mice exhibit a similar expansion of macrophages post-CIOA indicating that the inflammatory phase is not sex-specific. Finally, we find day 7 post-DMM is also characterized by increases in neutrophil, monocyte, and macrophage sub-population numbers. These results support a role for macrophages in early stages of OA through driving the inflammatory phase. Further investigation may elucidate potential targets for the prevention or attenuation of OA-associated cartilage damage.

## INTRODUCTION

Inflammation is a key factor in the development of osteoarthritis (OA). Although previously considered a disease of “wear and tear,” there is increasing evidence that OA patients exhibit synovitis (1-5). Increased cytokine levels in both synovial fluid and serum of early and stage OA suggest there is local and systemic inflammation (6-12). In fact, increased inflammation has been shown to correlate with worsened pain and cartilage damage (5). In post-traumatic OA (PTOA), where patients develop OA after a joint injury, synovitis has been shown to precede cartilage damage (13, 14). These patients may exhibit systemic inflammation even after corrective surgery (15). Macrophage infiltration into the joint synovium is often observed along with synovitis.

Macrophages are plastic immune cells that inhabit nearly every tissue in steady state and are capable of adapting to their residence. In response to signals in the local environment, macrophages reprogram their own regulatory landscape to perform tissue homeostatic functions (16, 17). In the naïve joint, the majority of synovial macrophages are self-maintained and radio-resistant (18). Some of these cells form a physical barrier as part of the synovial lining (19). Similar to other tissues, tissue-resident macrophages in the synovium are largely independent from circulation with minimal contribution from monocyte-derived macrophages (20-22). Macrophages are likely to be drivers of synovial inflammation through the secretion of inflammatory cytokines, extracellular matrix destruction, and recruitment of other immune cells, such as neutrophils (23, 24). Macrophages may also be directly responsible for cartilage damage as demonstrated through *in vitro* models (25, 26). The process by which macrophages are activated and transition from inflammation to overt symptoms of OA is not well understood.

Several mouse models exist that provide the opportunity to investigate experimental OA. Most of these models reflect PTOA since they are induced by injury, surgery, or enzyme (27-30). Collagenase-Induced OA (CIOA) is one such model that exhibits a distinct inflammatory phase preceding the establishment of overt joint damage (30). Moreover, macrophages have been previously implicated to play a role in cartilage damage in this model (31, 32). Thus, CIOA has the potential to reflect the role of inflammation in the development of OA in patients. Since CIOA entails a single injection it is relatively easy to perform on multiple mice in parallel while maintaining accurate and reproducible results (33-35). An alternative model, destabilization of the medial meniscus (DMM), involves micro-surgery performed by a trained mouse technician and is known to only produce reliable results in males (36). DMM is often preferred for modelling more progressive disease and has been used to investigate pain in experimental OA (37-39). Thus, CIOA and DMM are complementary models for studying the role of macrophages in post-traumatic OA.

In this study, we characterize macrophages in the knee joint during the development of experimental OA using both CIOA and DMM mouse models. We quantify immune cell infiltration and the proportion of macrophage sub-populations over the course of CIOA. By transcriptional profiling of synovial macrophages, we assess the function of different sub-populations in response to CIOA. We also compare immune cells numbers between male and female mice to determine whether there are sex effects. Finally, we investigate macrophages in the inflammatory phase of DMM. Collectively, our results indicate that macrophage are drivers of the inflammation that precedes cartilage damage in OA.

## RESULTS

### Infiltration of immune cells into the knee joint peaks on Day 3 post-CIOA

In CIOA, mice develop cartilage damage starting at 4 weeks (28 days) and exhibit established disease by 8 weeks (56 days) (30). To further investigate levels of inflammation in early OA, we assessed cellular infiltrates into the knee synovium of male mice that were untreated (day 0) and days 3, 7, 11, and 28 after collagenase was injected in both knees. We then processed joints for fluorescence-activated cell sorting and used a novel gating strategy to identify neutrophils, dendritic cells, monocytes, and macrophages from synovial tissue (**Figure 1A**). We found that all these inflammatory cells reached maximum numbers at day 3 although a high level of variability was observed across mice (**Figure 1B-E)**. Neutrophil and monocyte infiltration decreased rapidly, returning to steady-state levels by day7. On the other hand, macrophages, defined as CD11B+CD64+ cells were significantly increased on day 3 by approximately 8-fold (p=0.0248) and maintained an expanded population until at least day11. To confirm that macrophage infiltration was a result of the development of OA and not a response to the injection, we compared the knee of a mouse injected with collagenase with that of a mouse injected with PBS (**Supp. Figure 1A**). While macrophages in the collagenase-injected knees were significantly increased compared to naïve, PBS-injected knees were not (**Supp. Figure 1B**). Furthermore, the contralateral knee of the collagenase-injected mouse exhibited significantly fewer macrophages (**Supp. Figure 1B**). We observed similar effects on neutrophil numbers (**Supp. Figure 1B**). In summary, the peak of immune cell infiltration occurs around day 3 post-CIOA with a large increase in macrophage number.

**Figure 1.**
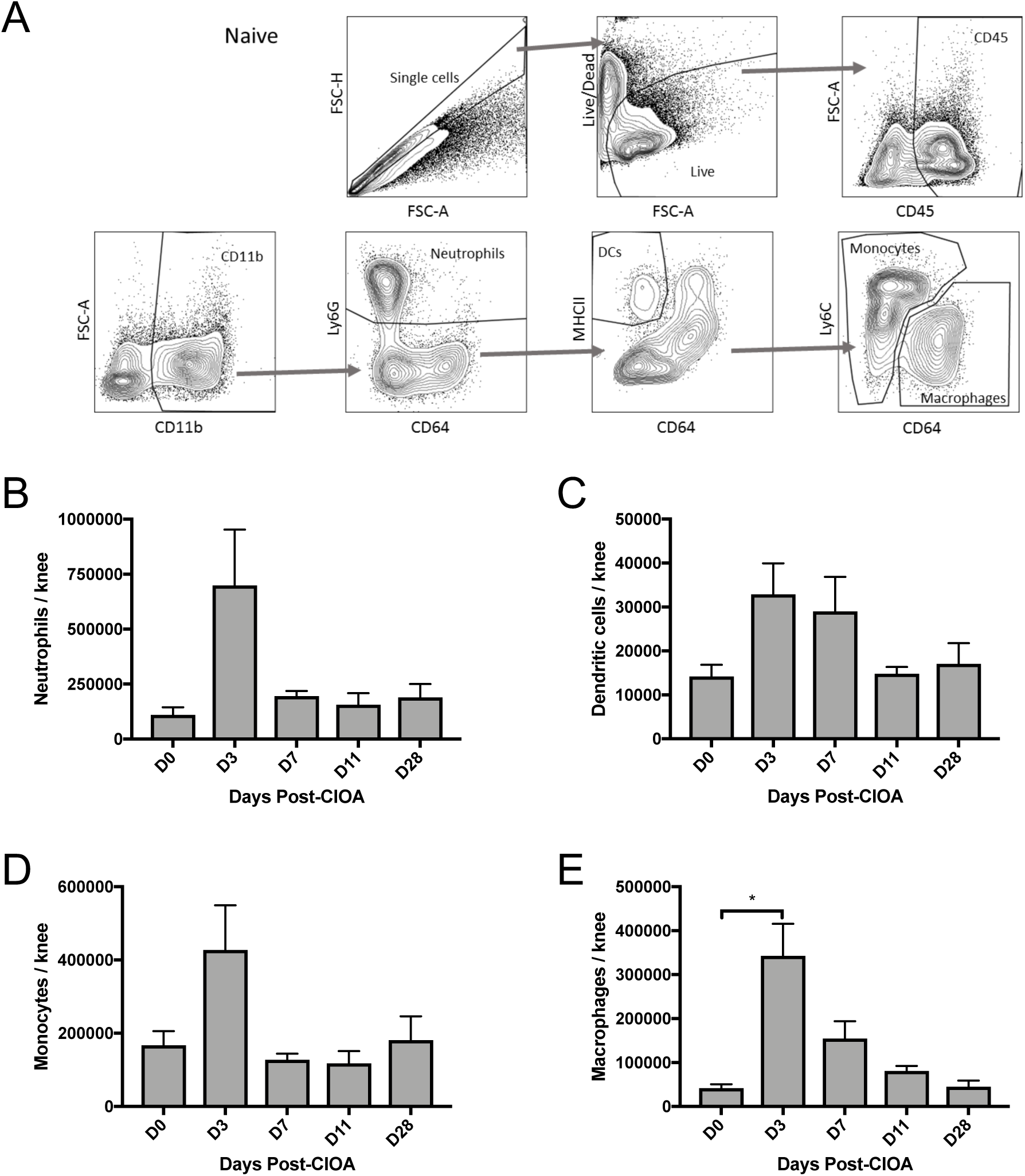
Immune cell infiltration in CIOA. (A) Gating strategy to identify neutrophils (Ly6G^+^CD64^-^), dendritic cells (DCs, MHCII^+^CD64^-^), monocytes (Ly6c^low-high^CD64^-^), and macrophages (CD64^+^) from CD45^+^CD11B^+^ synovial cells in the knee joints. (B) Quantification of neutrophils in the synovium per mouse with 2 knees injected with collagenase at days 0, 3, 7, 11, and 28 post-CIOA. (C) Quantification of DCs in the synovium per mouse with 2 knees injected with collagenase at days 0, 3, 7, 11, and 28 post-CIOA. (D) Quantification of monocytes in the synovium per mouse with 2 knees injected with collagenase at days 0, 3, 7, 11, and 28 post-CIOA. (E) Quantification of macrophages in the synovium per mouse with 2 knees injected with collagenase at days 0, 3, 7, 11, and 28 post-CIOA. N=3-4 mice per time point. *indicates p<0.05.

### The composition of the synovial macrophage compartment in the knee joint changes over the course of CIOA

Previous studies have demonstrated the heterogeneity of the macrophage compartment within tissues, such as lung and heart, even at steady-state (40, 41). In the synovial tissue, prior work has suggested that MHCII surface marker expression distinguished tissue-resident (MHCII-) from monocyte-derived (MHCII+) macrophages (18). Further, a recent study demonstrated that synovial lining macrophages are CX3CR1+ (19). Thus, we distinguished for sub-populations of synovial macrophages and quantified them over the course of CIOA: CX3CR1+MHCII+ (MA), CX3CR1+MHCII+ (MB), CX3CR1-MHCII- (MC), and CX3CR1-MHCII+ (MD) (**Figure 2A**). We found that all four macrophage sub-populations exhibited a significant increase in numbers on day 3 post-CIOA (**Figure 2B-E**, p<0.05). The expansion of MA and MB macrophages (∼14- and 11-fold) was greater than MC and MD (∼6- and 5-fold) between day 3 and day 0. The MA sub-population, which likely represents synovial lining macrophages accounted for a greater proportion of the macrophage compartment at peak inflammation on day 3 but returned to steady-state levels by day 28 (**Figure 2F, Supp. Figure 2**). However, the proportion of MB macrophages, which we believe to be monocyte-derived, is still increased at day 28 (**Figure 2F, Supp. Figure 2**). Taken together, these data support an expansion of the synovial macrophage compartment that leads to a disproportionate increase in synovial lining and monocyte-derived macrophages.

**Figure 2.**
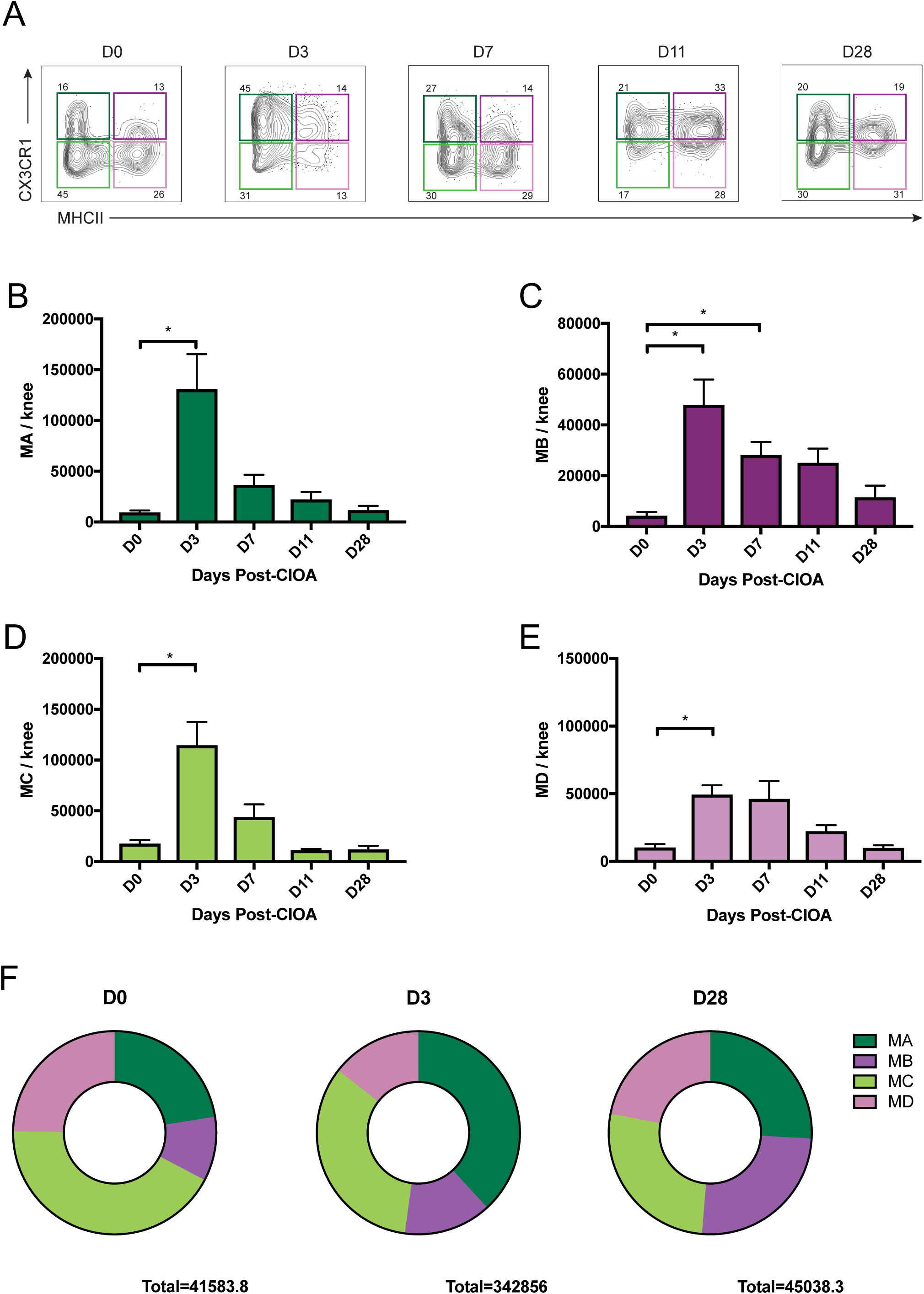
Macrophage sub-population numbers over the course of CIOA. (A) Representative gating of macrophage sub-populations from CD45^+^CD11B^+^ CD64^+^ synovial cells in the knee joints at days 0, 3, 7, 11, and 28 post-CIOA: CX3CR1+MHCII+ (MA), CX3CR1+MHCII+ (MB), CX3CR1-MHCII- (MC), and CX3CR1-MHCII+ (MD). (B) Quantification of MA macrophages in the synovium per mouse with 2 knees injected with collagenase at days 0, 3, 7, 11, and 28 post-CIOA. (C) Quantification of MB macrophages in the synovium per mouse with 2 knees injected with collagenase at days 0, 3, 7, 11, and 28 post-CIOA. (D) Quantification of MC macrophages in the synovium per mouse with 2 knees injected with collagenase at days 0, 3, 7, 11, and 28 post-CIOA. (E) Quantification of MD macrophages in the synovium per mouse with 2 knees injected with collagenase at days 0, 3, 7, 11, and 28 post-CIOA. (F) Mean proportion of macrophage sub-populations in the knee synovium at days 0, 3, and 28 post-CIOA. N=3-4 mice per time point. *indicates p<0.05.

### Synovial macrophages exhibit a peak inflammatory response in CIOA

In order to investigate the function of synovial macrophages during the development of OA, we performed RNA-seq on the four macrophage sub-populations isolated from the mice above on days 0, 3, 7, 11, and 28 post CIOA. We found that each macrophage sub-population exhibited distinct gene expression at day 0 but their transcriptional profiles appeared to converge at peak inflammation on day 3 (**Figure 3A, Supp Figure 3A-B**). By resolution of inflammation on day 28, the transcriptional profile of each sub-population resembled steady-state. To determine the similarity in the response of the different macrophage sub-populations in the inflammatory phase of CIOA, we compared the differentially expressed genes between days 0 and 3 (**Supplemental Figure 3C**). We found that the highest overlap between any sub-population is that between all four sub-populations for both up- (106) and down- (111) regulated genes (**Figure 3B**). Notably, MA macrophages exhibit a high number of unique differentially expressed genes, suggesting that this sub-population undergoes a distinct response in CIOA.

**Figure 3.**
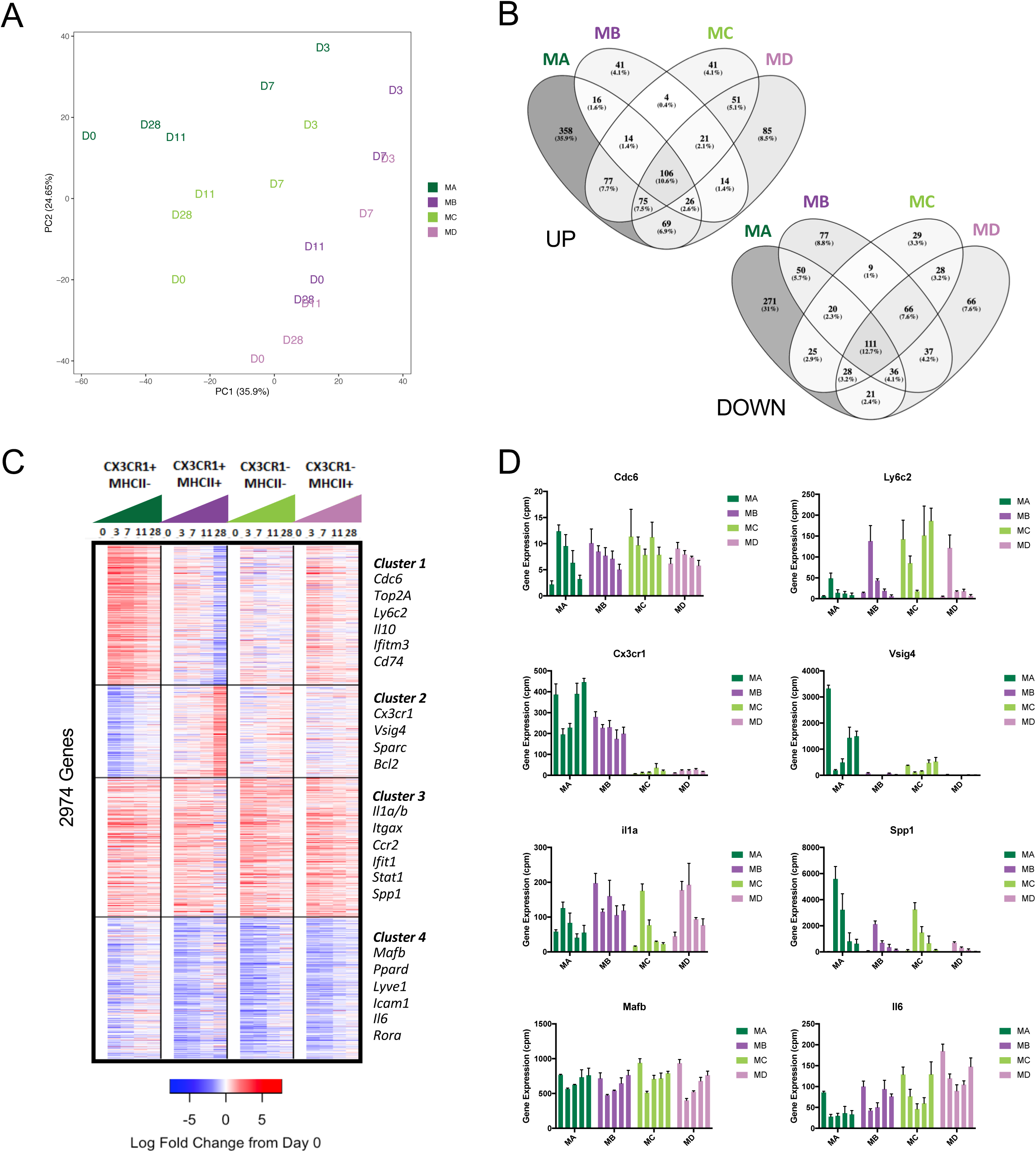
Transcriptional profiles of macrophage sub-populations over the course of CIOA. (A) Principal component analysis (PCA) of mean gene expression as measured by RNA-seq from CX3CR1+MHCII+ (MA), CX3CR1+MHCII+ (MB), CX3CR1-MHCII- (MC), and CX3CR1-MHCII+ (MD) on days 0, 3, 7, 11, and 28 post-CIOA. (B) Overlap of differentially expressed genes between naive (day 0) and peak inflammation (day 3) of CIOA in each macrophage sub-population. [left=increased on day 3; right = decreased on day 3] (C) k-means clustering (k=4) of the mean fold-change of 2974 differentially expressed genes across the CIOA time course in at least one macrophage sub-populations. Example genes in each cluster are indicated on the right. (D) Expression of individual genes in each macrophage sub-population over the course of CIOA. N=3-4 mice per time point.

To further define the overall patterns of gene expression, we clustered 2974 genes that were differentially expressed across the CIOA time course into four clusters by their fold-change in gene expression relative to day 0 using k-means (**Figure 3C**). Clusters 1 and 2 represent genes that are primarily up- and down-regulated in the MA macrophages, respectively, and are consistent with the results above. The up-regulated genes in cluster 1 included many associated with *cell cycle*, such as Cdc6 and Top2a, and monocytic origin (42), such as Ly6c2 and CD52 **(Supp. Table 1)**, suggesting that the expansion of MA sub-population may be a result of proliferation as well as infiltrating monocytes. In addition, several immune-related GO processes including response to interferon-gamma and MHCII antigen presentation, such as Il10, Ifitm3, and CD74, were enriched in cluster 1 **(Supp. Table 1)**. In contrast, cluster 2 contains genes previously identified as part of the synovial lining phenotype – such as Cx3cr1, Sparc, and Vsig4, indicating that MA cells are losing their distinct identity (19). These genes were also connected to “NUPR1+” myeloid cells from synovial biopsies of OA and rheumatoid arthritis (RA) patients which were inversely correlated with tissue inflammation (43) (**Supp. Figure 3D**). Clusters 3 and 4 represent genes exhibit similar patterns of gene expression across macrophage sub-populations. In particular, cluster 3 reflects the increase in specific inflammatory genes in all synovial macrophages including those associated with regulation of IL-1B/IL-6, response to interferon beta, and additional monocyte genes (Ccr2, Itgax, and Irf7) **(Supp. Table 1)**. Interestingly, this cluster also included gene implicated in osteoclast differentiation, such as Spp1, Acp5, and Ctsk (44, 45). Finally, cluster 4 comprised genes associated with cell adhesion and homeostasis, such as Icam1, Ppard, and Lyve1 **(Supp. Table 1)**: the down-regulation of these genes may indicate that that synovial macrophages are prioritizing inflammatory response genes over steady-state functions. The transcription factors, MafB, which is associated with macrophage maturity was also found in this cluster (46). Although genes associated with both “IFN-activated” and “IL1B+ pro-inflammatory” myeloid cells we up-regulated in synovial biopsies from RA patients, cluster 3 was associated with the former and cluster 4 was associated with the latter (**Supp. Figure 3D**). This may indicate that OA-associated inflammation is distinct from that observed in RA (43). Therefore, these results suggest that synovial macrophages increase expression of genes associated with OA-specific inflammation and monocyte infiltration, while decreasing expression of genes associated with the tissue-resident phenotype.

### Female mice exhibit a similar inflammatory response post-CIOA

Typically, male mice are used for experimental OA since female mice have been shown to exhibit distinct immune responses in many models, including DMM (47). However, we observed that the knee joints of female mice exhibit a comparable level of articular cartilage damage to male mice at 8 weeks post-CIOA (**Figure 4A**). To determine whether female mice respond similarly to CIOA, we assessed immune cell infiltration over a similar time course to the male mice. We found that total macrophage number in female mice was significantly increased at day 3 post-CIOA (p= 0.041) and then progressively decreased to steady-state levels by day 28 (**Figure 4B**). The expansion of macrophage in females was comparable to that observed in male, although the fold-change at day 3 was somewhat lower (∼5-fold vs ∼8-fold) (**Figure 4C**). Other immune cell populations also observed similar trends between the sexes with neutrophil numbers significantly increased at day 3 (p=0.033) (**Supp. Figure 4A-C**). Moreover, we found that each macrophage sub-population in female mice peaked at day3 post-CIOA and returned to approximately steady-state by day 28 (**Figure 4D**). Compared to male mice, female mice exhibit an increased proportion of MC and MD sub-populations at day 0 and end the time-course with a larger proportion of MD rather than MB as observed in males (**Supp. Figure 4D**). Since MC and MD represent the CX3CR1-sub-populations, this result may be related to differing CX3CR1 signaling in females as observed previously in microglia, the brain-resident macrophages (48). Overall, our studies suggest that female mice exhibit a similar inflammatory response in experimental OA.

**Figure 4.**
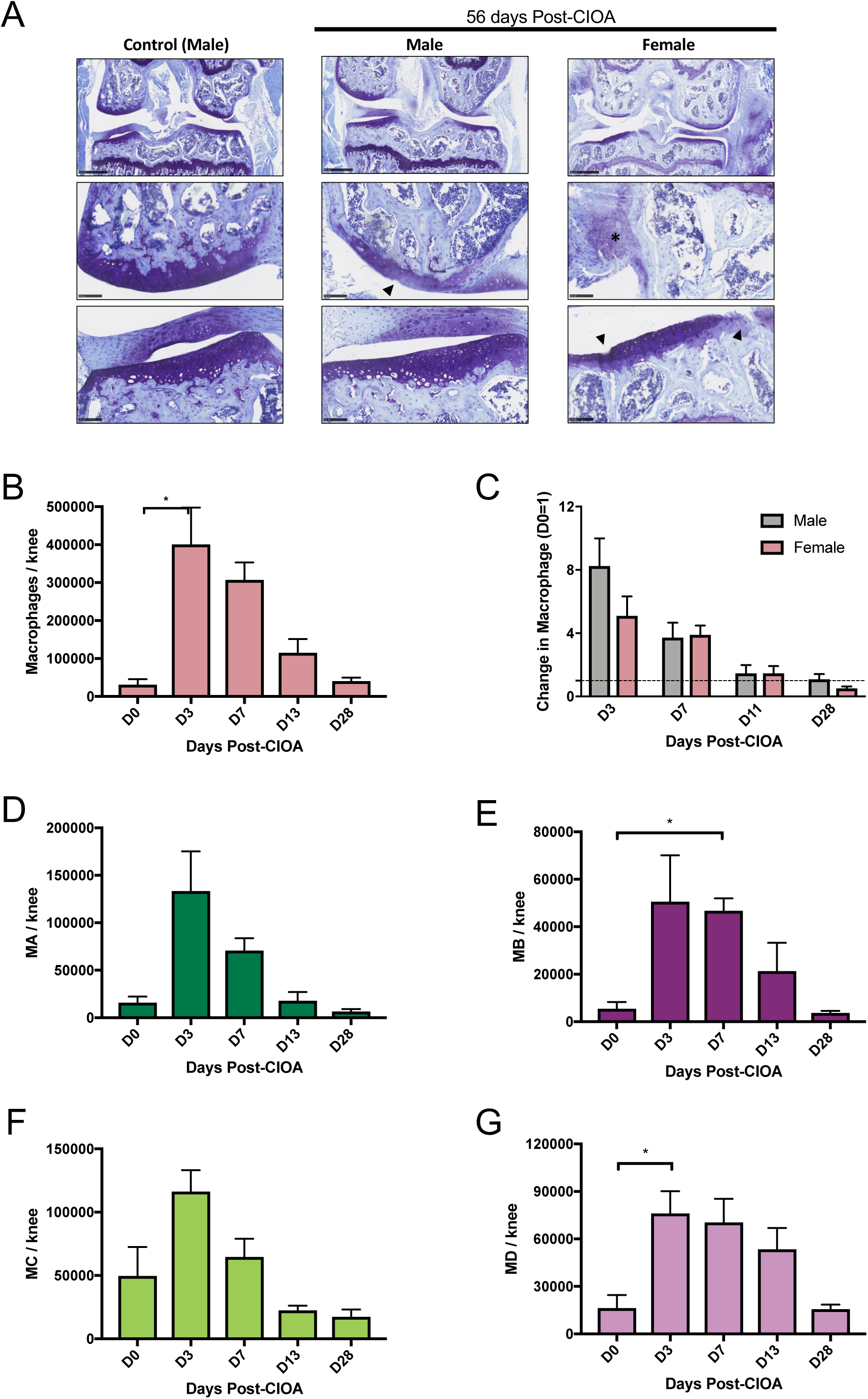
Comparison of macrophage sub-populations over the course of CIOA in female mice. (A) Frontal histology of late stage OA at 8 weeks in males and females. **\*** indicates chondrophyte, Δ indicates articular cartilage damage. (B) Quantification of total macrophages in the synovium per female mouse with 2 knees injected with collagenase at days 0, 3, 7, 13, and 28 post-CIOA. (C) Fold-change in total macrophage number relative to day 0 in the synovium per female mouse with 2 knees injected with collagenase at days 0, 3, 7, 13, and 28 post-CIOA. (D) Quantification of MA macrophages in the synovium per female mouse with 2 knees injected with collagenase at days 0, 3, 7, 13, and 28 post-CIOA. (E) Quantification of MB macrophages in the synovium per female mouse with 2 knees injected with collagenase at days 0, 3, 7, 13, and 28 post-CIOA. (F) Quantification of MC macrophages in the synovium per female mouse with 2 knees injected with collagenase at days 0, 3, 7, 13, and 28 post-CIOA. (G) Quantification of MD macrophages in the synovium per female mouse with 2 knees injected with collagenase at days 0, 3, 7, 13, and 28 post-CIOA. N=3-4 mice per time point. *indicates p<0.05.

### Macrophage infiltration also defines response to DMM

DMM is a widely used mouse model for post-traumatic OA that is generally considered to be less inflammatory than CIOA (36). However, prior work suggests that mice experience synovial inflammation for 2 weeks post-DMM (49). Thus, we assessed immune cell infiltration in male mice at day 7 post-DMM using the same gating strategy described above (**Figure 1A**). We found that neutrophils, monocytes, and macrophages, but not DCs, were significantly increased compared to both untreated knees from naïve mice (p< 0.05) (**Figure 5A-D, Supp. Figure 5A)**. Moreover, the synovium of the contralateral knee also exhibited significant increases in neutrophil and monocyte infiltration suggesting that there is some systemic inflammation resulting from DMM surgery. On the other hand, macrophages, which expand more than 6-fold in the DMM knee, are not significantly different in the contralateral joint. Similarly, each of the four macrophage sub-populations are increased in the DMM knee compared to naïve or contralateral (**Figure 5E-F**). As in CIOA males, the CX3CR1+ macrophage, both the MA synovial lining and MB monocyte-derived, exhibited the greatest increase in numbers (**Supp. Figure 5B**). Thus, the macrophage-driven inflammatory phase may be a general feature of experimental OA.

**Figure 5.**
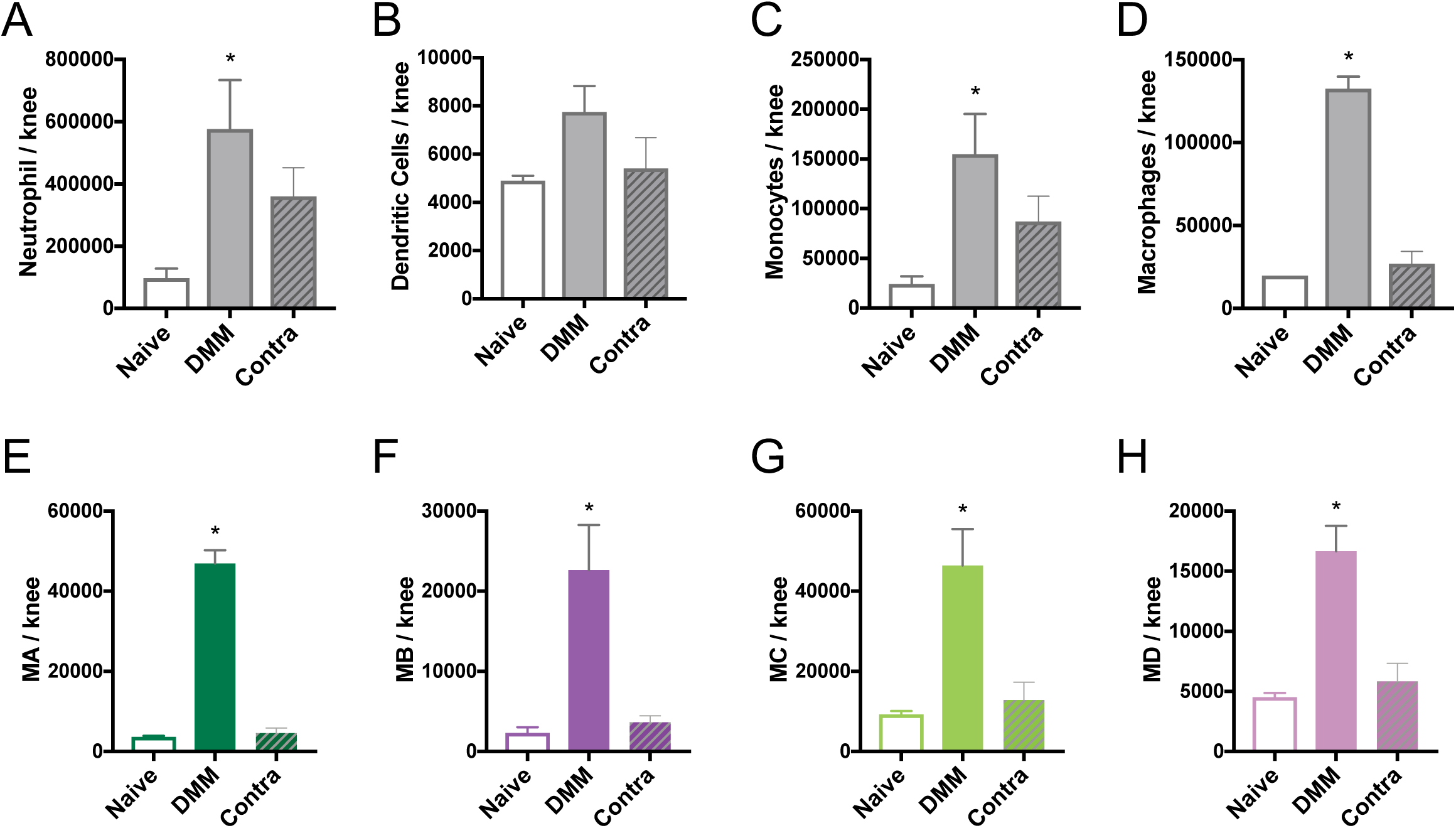
Immune cell infiltration in DMM. (A) Quantification of neutrophils in the synovium per knee with no treatment (naïve), DMM surgery (DMM), and opposite from DMM surgery (contralateral) at days 7 post-surgery. (B) Quantification of DCs in the synovium per knee with no treatment (naïve), DMM surgery (DMM), and opposite from DMM surgery (contralateral) at days 7 post-surgery. (C) Quantification of monocytes in the synovium per knee with no treatment (naïve), DMM surgery (DMM), and opposite from DMM surgery (contralateral) at days 7 post-surgery. (D) Quantification of total macrophages in the synovium per knee with no treatment (naïve), DMM surgery (DMM), and opposite from DMM surgery (contralateral) at days 7 post-surgery. (E) Quantification of MA macrophages in the synovium per knee with no treatment (naïve), DMM surgery (DMM), and opposite from DMM surgery (contralateral) at days 7 post-surgery. (F) Quantification of MB macrophages in the synovium per knee with no treatment (naïve), DMM surgery (DMM), and opposite from DMM surgery (contralateral) at days 7 post-surgery. (G) Quantification of MC macrophages in the synovium per knee with no treatment (naïve), DMM surgery (DMM), and opposite from DMM surgery (contralateral) at days 7 post-surgery. (H) Quantification of MD macrophages in the synovium per knee with no treatment (naïve), DMM surgery (DMM), and opposite from DMM surgery (contralateral) at days 7 post-surgery. N=4 mice per group. *indicates p<0.05.

## DISCUSSION

Macrophages are critical cells in both steady-state homeostasis and immune response. Hence, they may fulfill a variety of functions in the development of OA. Here, we demonstrate that multiple sub-populations of macrophages are expanded in the knees of experimental OA using two distinct and well-established mouse models: CIOA and DMM (30, 36). As demonstrated by transcriptional profiling, these synovial macrophages alter their function in response to joint injury. A large number of the transcriptional changes are shared across all macrophage sub-populations. In general, these changes reflect those typical of immune response: an up-regulation of inflammatory genes coinciding with a down-regulation of homeostatic genes. Similar responses are observed in other tissues, such as in alveolar macrophages during pulmonary fibrosis or Kupffer cells in liver disease (40, 50). A more in depth analysis of the genes expressed dynamically across the CIOA time course in each sub-population may reveal more specific processes. For example, genes primarily down-regulated in the MA sub-population are associated with the tissue-resident phenotype. Since we propose that MA macrophages form the synovial lining, this sub-population expresses these genes at the highest level at steady-state. MB exhibit an increase in these genes much later around day 28 post-CIOA, which may reflect the transition of these cells from newly monocyte-derived macrophages into tissue-resident macrophages.

The infiltration of monocytes from circulation may be largely responsible for the expansion of synovial macrophages. In steady-state, the minority of macrophages in the joint are recognizable as monocyte-derived by MHCII+ surface marker expression, while the remaining are likely to be long-lived embryonically derived cells (18). As in other tissues, when the macrophage compartment is experimentally depleted, circulating cells are capable of extravasating and differentiating into macrophages. These steady-state monocyte-derived macrophages may be indistinguishable, even on a transcriptional and epigenomic level, from true tissue-resident cells (17, 51). As plastic cells, macrophages are capable of adapting their regulatory programming in response to signal in the local environment (52). However, in the case of an immune challenge, such as the initiation of OA, the local environment is perturbed. Infiltrating inflammatory monocytes may not follow the typical path to differentiate into tissue-resident macrophages, instead maintaining expression of monocyte-associated genes as we observed. It is also possible that synovial macrophages may proliferate to increase their numbers in OA. In support of this alternative, our data demonstrated an enrichment of cell cycle gene specifically in the cluster up-regulated in MA macrophages. Together, these results may support the previously proposed model where proliferating macrophages and infiltrating monocytes compete to fill the macrophage niche (53).

The data presented here reflects macrophage involvement in the development of experimental OA. Since we observe similar results in both sexes in CIOA and in the additional model of DMM, we propose that the role of macrophages is conserved across different OA etiologies. Macrophages have been similarly implicated in the development of disease in OA patients. However, it is difficult to obtain data on the early stages of preclinical OA. Typically, samples are obtained from patients with established disease or even terminal disease. In our prior work, we compared synovial biopsies from RA patients with synovial tissues obtained from joint replacements on OA patients (54). Thus, the transcriptional profile of macrophages in these samples are unlikely to match our observation in experimental OA. However, we did observe that a subset of the OA patient exhibited a transcriptional signature of inflammation comparable to RA. This observation was shown independently by another group that classified two subtypes of macrophages in OA (55). Furthermore, AMP characterized myeloid sub-populations from joint replacements and compared the transcriptional profile of macrophages from OA and RA patients (43). In particular, their OA macrophages exhibited increased expression of genes that were down-regulated in our data. Since these samples reflect very late stage disease, these genes may have already returned to their steady-state levels as seen by day 28 in CIOA. Mouse models provide the opportunity to investigate the role of macrophages in the inflammatory phase of experimental OA.

Understanding the function of macrophages in the development of OA is likely to provide insight into prevention or suspension of disease. Because macrophages are responsive to their environment, they offer a readout of the immune state of the synovium. Our data suggest that changes to the macrophage transcriptional profile can predict the development of cartilage damage. Moreover, macrophages themselves may be an avenue for future therapeutics to alleviate the inflammation that gives way to disease. Additional investigation into the underlying regulatory factors driving macrophage phenotype in experimental may shed light on possible targets. Comparison of these data with human samples, preferably during early stage disease, such as at the time of ACL repair in OA susceptible patients, can further support these studies.

## METHODS

### Mice

The genetic background of mice in all studies was C57Bl/6. Mice were housed in barrier and specific pathogen-free facilities at Northwestern University or Rush University Medical College and subject to continual health monitoring. Male mice aged 8-12 weeks were used unless otherwise stated.

### CIOA injection

The CIOA model was performed as previously described (30, 33-35). Briefly, arthritis is induced by an intra-articular injection of collagenase type VII (LPS-free) into the synovial space of the knee. Mice were anesthetized via isofluorane and the procedure was performed under continual supply of anesthetic via nose-cone. Once fully anesthetized, the injection site was prepared by hair removal and washing the site with 70% ethanol. With the animal’s knee bent at a 90 degree angle, a very small incision (<5mm in length) is made directly over the patella to expose the joint. Freshly prepared sterile collagenase type VII is diluted in sterile PBS pH 7.4 to a concentration of 1.66U/*μ*L. Delivery of the collagenase into the joint is performed using a Hamilton Syringe with a 30ga syringe needle to ensure precision. The needle point is carefully inserted under the patellar tendon into the synovial space, and 6*μ*l of collagenase solution is injected. The knee is then manually articulated to ensure distribution of the collagenase solution within the synovium. For the day 3 to 28 time course, both knees were injected with collagenase and then pooled or used for different studies. To assess the effect on the contralateral knees, collagenase was injected into left knees and the right knees were processed in parallel on day 3. As an additional control, we followed the same procedure with a PBS-only injection.

### DMM surgery

The DMM model was performed as described previously (38, 39). Briefly, mice are anaesthetized by Ketamine/Xylazine (100mg/kg, 5mg/kg respectively, i.p.), the right hind limb is shaved and swabbed with 70% ethanol, the animal positioned on a dissecting microscope and the leg draped. A medial para-patella arthrotomy is performed and the patella luxated laterally. The anterior fat pad is dissected to expose the anterior medial meniscotibial ligament, which is elevated and severed using curved dissecting forceps 10. Complete severance of the ligament is confirmed visually by manually displacing the medial meniscus. The patella is repositioned, the knee is flushed with saline and the incision closed in 3 layers-simple continuous 8/0 vicryl in the joint capsule, simple continuous 8/0 vicryl subcutaneously and tissue glue for the skin. To assess the effect on the contralateral knee, surgery was performed on right knees and left knees were processed in parallel.

### Processing of synovial tissue

Mice were euthanized and one or both knees was dissected, extraneous muscle tissue resected, and joints processed separately unless otherwise stated. For histology, knees were fixed, decalcified, and cut frontally to assess articular cartilage damage. For Fluorescence-Activated Cell Sorting (FACS), bone marrow is removed along with medial section of the leg bones and knees were perfused with 1mL of digestion buffer (dispase II, collagenase D, and DNase I in HBSS). Synovium was exposed by cutting through the patellar tendon into the joint space. The knees are incubated in 2mL of additional digestion buffer for one hour at 37°C in an orbital shaker. Post-digestion, the knees and digestion buffer are filtered using a 40μm nylon mesh filter and remaining synovial tissue is agitated against the mesh for optimal cell recovery. Erythrocyte lysis is performed using 1x PharmLyse, then the cells are stained for flow cytometry. Cells are incubated with a live/dead stain, FC Block, and fluorochrome-conjugated antibodies for flow cytometry analysis (PerCPCy5.5 CD11b; eFluor450 MHC11; Alexa 647 CX3CR1; AF700 CD45; APCCy7 Ly6C; PE CD64; PECF594 Siglec F, CD4, CD8, CD19, NK1.1; PECy7 Ly6G). Data collection and analysis is performed using BD FACSDiva Software and FlowJo Software. Differences between cell numbers at different tie points or conditions was calculated by unpaired t-test with Welch correction for unequal variance.

### RNA-sequencing

Macrophage sub-populations isolated by FACS were re-suspended in Picopure RNA Isolation (ThermoFisher) lysis buffer and RNA was extracted as per manufacturer’s instructions. Library prep was performed using full-length SMART-seq v4 Ultra Low Input Kit (Clontech) and sequenced on Illumina NextSeq. The resulting bcl sequencing files were demultiplexed using bcl2fastq and processed using a custom pipeline. Briefly, the reads were trimmed using Trimmomatic version 0.36 (56), aligned to the mm10 genome with tophat 2.1.0 (57), and mapped to gene exons by HTseq (unstranded) (58) to generate a table of gene expression counts. To account for differing read depth across samples, we converted raw counts to counts per million (cpm). Expressed genes were defined by filtering for genes with expression greater than 7 cpm in at least 4 samples, leaving 10581 genes. Differential expression between day 0 and day 3 post-CIOA were calculated by DEseq (59) and genes were considered differential if the estimated log-fold change was greater than 1 or less -1 and the adjusted p-value was less 0.05. Differentially expressed genes across the CIOA time course were defined as those with at least a log fold-change of between the mean of any 2 time-points. Time course RNA-seq was visualized as the fold-change between the mean of each post-CIOA time point and day 0. Enrichment of marker genes from Zhang et al (43) were calculated using the hypergeometric distribution on the proportion of genes in each of the four k-means clusters compared to total expressed genes.

## FIGURE LEGENDS

**Supplemental Figure 1. Immune cell infiltration in CIOA**. (A) Representative gating of neutrophils (Ly6G^+^CD64^-^), dendritic cells (DCs, MHCII^+^CD64^-^), monocytes (Ly6c^low-high^CD64^-^), and macrophages (CD64^+^) from CD45^+^CD11B^+^ synovial cells isolated from both knee joints of untreated mice (naïve), the left knee of mice injected with PBS (PBS-injected), and the left (collagenase-injected) and right (collagenase contralateral) knees of mice injected with collagenase at day 3 post-treatment. (B) Quantification of neutrophils (top), DCs (second from top), monocytes (second from bottom), and macrophages (bottom) in the synovium of naïve, PBS-injected, collagenase-injected, and collagenase contralateral synovium per knee at day 3. N=3 knees per group. *indicates p<0.05.

**Supplemental Figure 2. Macrophage sub-population numbers over the course of CIOA**. Mean proportion of macrophage sub-populations in the knee synovium at days 7 and 11 post-CIOA: CX3CR1+MHCII+ (MA), CX3CR1+MHCII+ (MB), CX3CR1-MHCII- (MC), and CX3CR1-MHCII+ (MD). N=3-4 mice per time point.

**Supplemental Figure 3. Transcriptional profiles of macrophage sub-populations over the course of CIOA**. (A) Pairwise Pearson correlation between mean gene expression as measured by RNA-seq from CX3CR1+MHCII+ (MA), CX3CR1+MHCII+ (MB), CX3CR1-MHCII- (MC), and CX3CR1-MHCII+ (MD) on days 0, 3, 7, 11, and 28 post-CIOA. (B) Principal component analysis (PCA) of gene expression as measured by RNA-seq from replicates of CX3CR1+MHCII+ (MA), CX3CR1+MHCII+ (MB), CX3CR1-MHCII- (MC), and CX3CR1-MHCII+ (MD) on days 0, 3, 7, 11, and 28 post-CIOA. (C) Volcano plots of differentially expressed genes between naive (day 0) and peak inflammation (day 3) of CIOA in each macrophage sub-population. (D) Enrichment in clusters 1:4 from **Figure 3C** of marker genes from myeloid cell sub-populations (SC-M1:4) in synovial tissue of OA and RA patients (43). White bar indicates the expected number of genes in each cluster given its size. * indicates p<0.05 N=3-4 mice per time point.

**Supplemental Figure 4. Comparison of macrophage sub-populations over the course of CIOA in female mice**. (A) Quantification of neutrophils in the synovium per female mouse with 2 knees injected with collagenase at days 0, 3, 7, 13, and 28 post-CIOA. (C) Quantification of DCs in the synovium per female mouse with 2 knees injected with collagenase at days 0, 3, 7, 13, and 28 post-CIOA. (D) Quantification of monocytes in the synovium per female mouse with 2 knees injected with collagenase at days 0, 3, 7, 13, and 28 post-CIOA. (D) Mean proportion of macrophage sub-populations in the knee synovium of female mice at days 0, 3, 7, 13, and 28 post-CIOA: CX3CR1+MHCII+ (MA), CX3CR1+MHCII+ (MB), CX3CR1-MHCII- (MC), and CX3CR1-MHCII+ (MD). N=3-4 mice per time point. *indicates p<0.05.

**Supplemental Figure 5. Immune cell infiltration in DMM**. (A) Representative gating of neutrophils (Ly6G^+^CD64^-^), dendritic cells (DCs, MHCII^+^CD64^-^), monocytes (Ly6c^low-high^CD64^-^), and macrophages (CD64^+^) from CD45^+^CD11B^+^ synovial cells isolated from the knees of untreated mice (naïve) and the treated (DMM) and opposite (contralateral) knees of mice on day 7 post-surgery. (B) Representative gating of macrophage sub-populations from CD45^+^CD11B^+^ CD64^+^ synovial cells in the knees of untreated mice (naïve) and the treated (DMM) and opposite (contralateral) knees of mice on day 7 post-surgery CX3CR1+MHCII+ (MA), CX3CR1+MHCII+ (MB), CX3CR1-MHCII- (MC), and CX3CR1-MHCII+ (MD).

