## Supplemental Figures 1-5 for "Macrophages drive the inflammatory phase in experimental osteoarthritis"

Supplemental Figure 1.

A

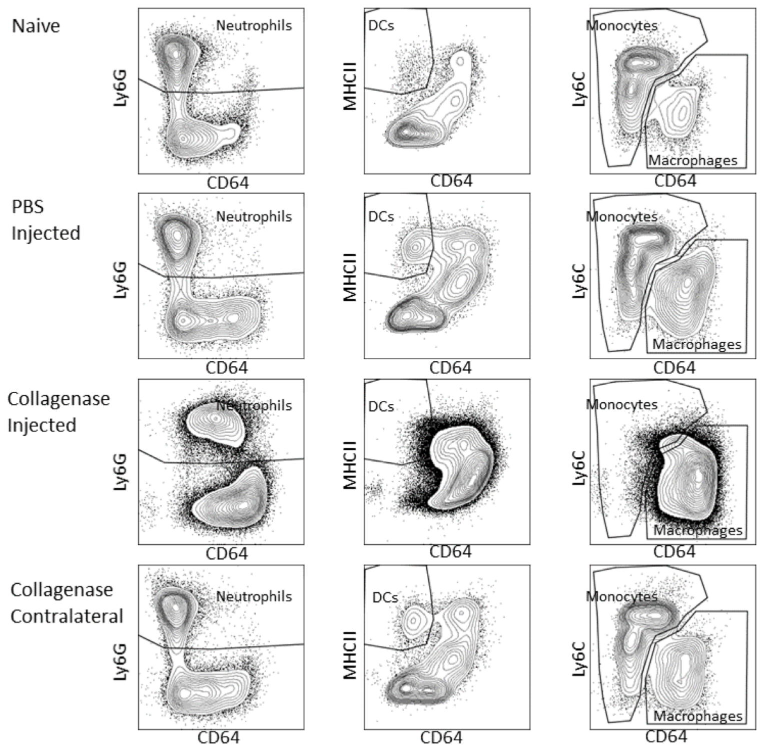


B

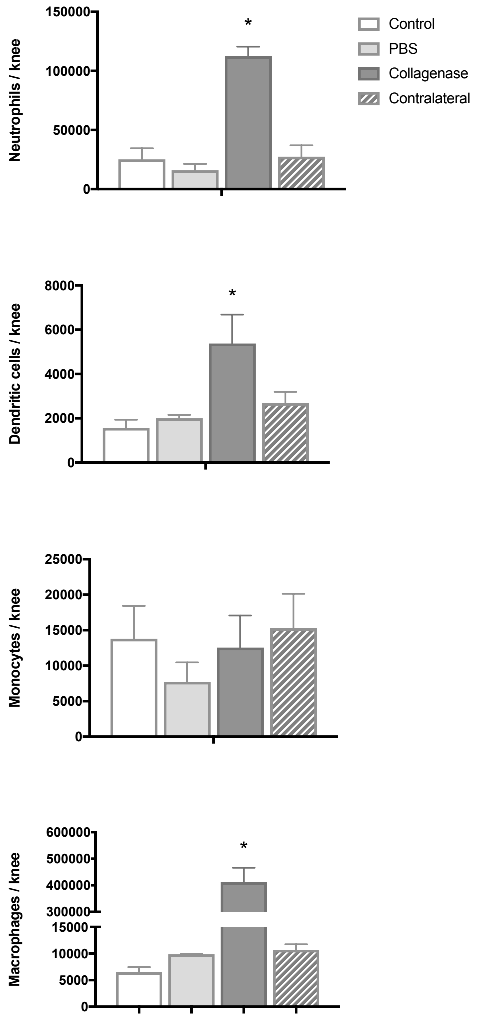

Supplemental Figure 2.

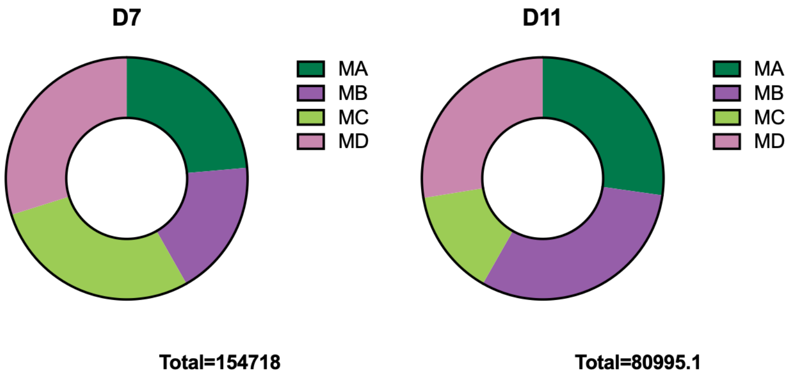

Supplemental Figure 3.

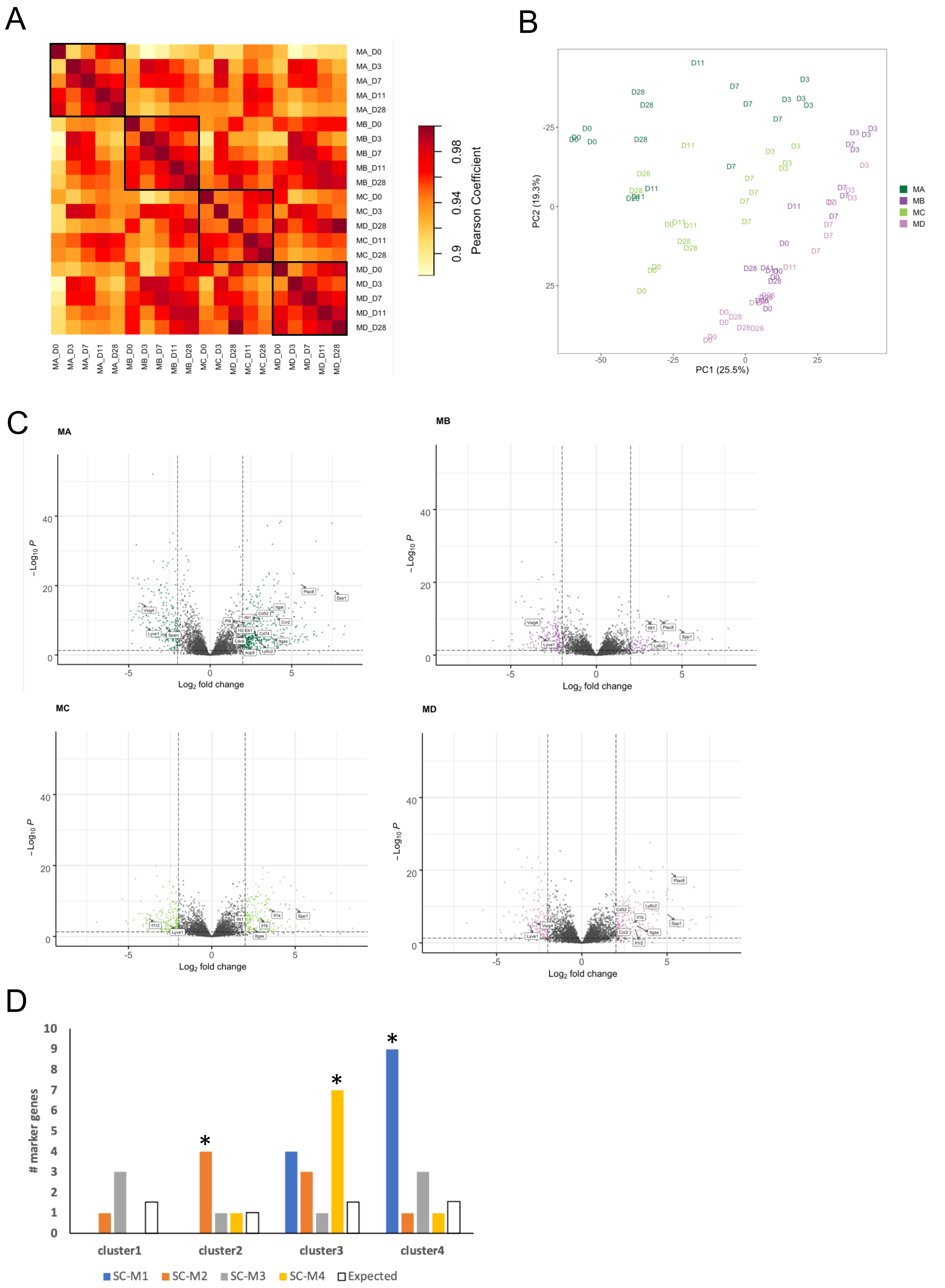

Supplemental Figure 4.

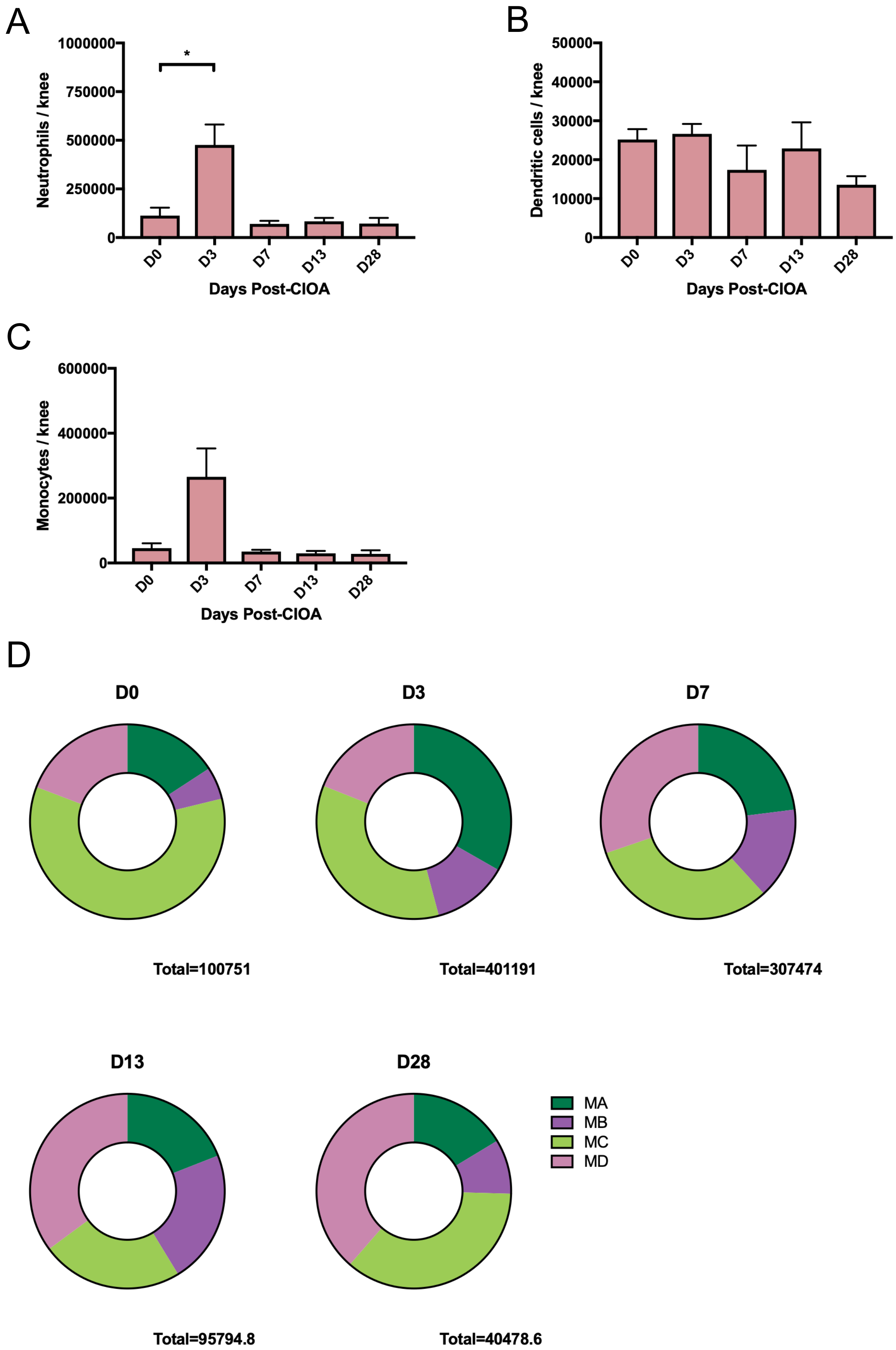

Supplemental Figure 5.

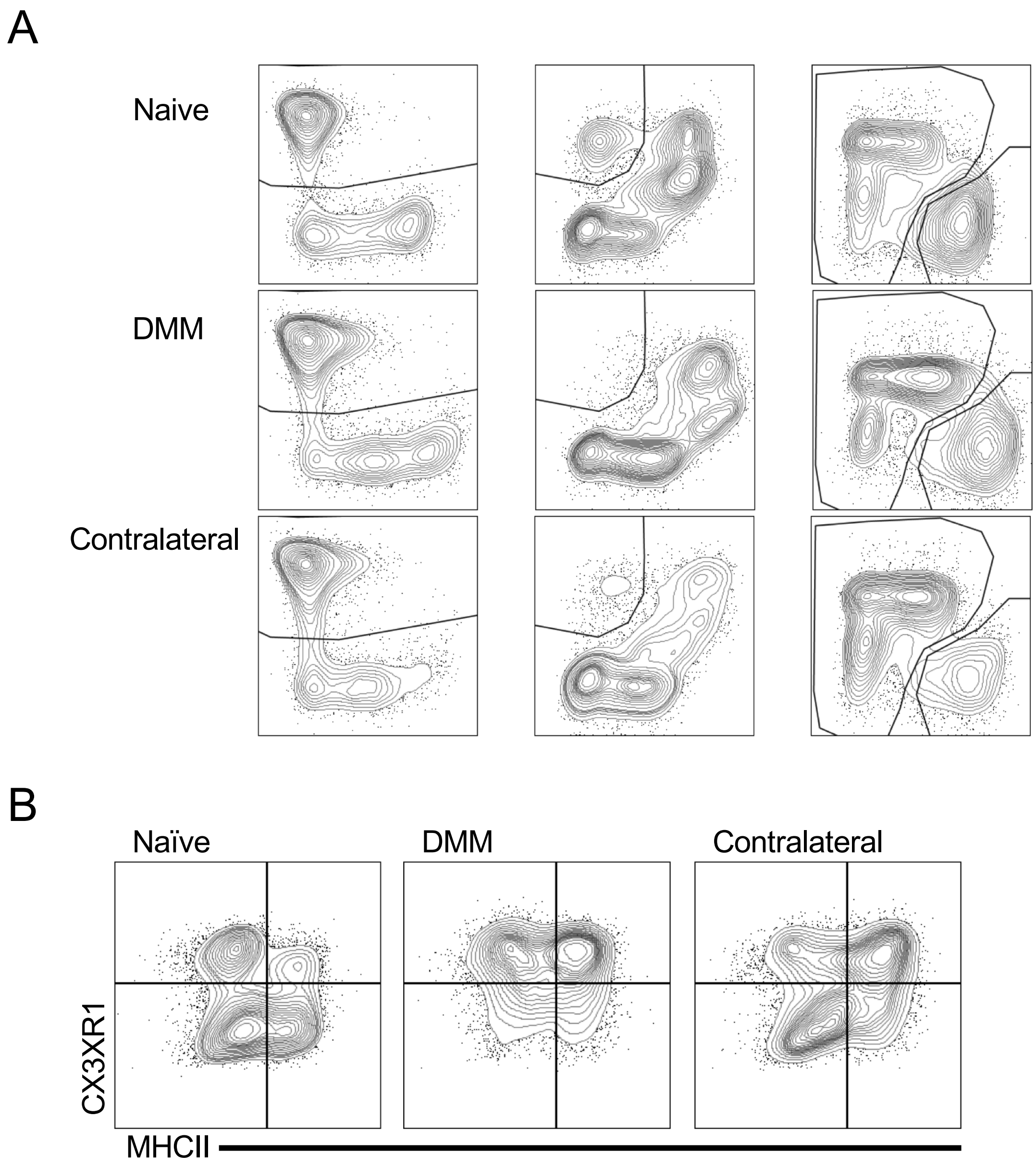
